# A rapidly evolving polybasic motif modulates bacterial detection by guanylate binding proteins

**DOI:** 10.1101/689554

**Authors:** Kristin M. Kohler, Miriam Kutsch, Anthony S. Piro, Graham Wallace, Jörn Coers, Matthew F. Barber

**Affiliations:** Institute of Ecology & Evolution, University of Oregon. Eugene, OR 97403; Department of Molecular Genetics and Microbiology, Duke University Medical Center., Durham, NC 22710; Department of Immunology, Duke University Medical Center. Durham, NC 22710; Department of Biology, University of Oregon. Eugene, OR 97403

## Abstract

Cell-autonomous immunity relies on the rapid detection of invasive pathogens by host proteins. Guanylate binding proteins (GBPs) have emerged as key mediators of vertebrate immune defense through their ability to recognize a diverse array of intracellular pathogens and pathogen-containing cellular compartments. Human and mouse GBPs have been shown to target distinct groups of microbes, although the molecular determinants of pathogen specificity remain unclear. We show that rapid diversification of a C-terminal polybasic motif (PBM) in primate GBPs controls recognition of the model cytosolic bacterial pathogen *Shigella flexneri*. By swapping this membrane-binding motif between primate GBP orthologs, we find that the ability to target *S. flexneri* has been enhanced and lost in specific lineages of New World monkeys. Single substitutions in rapidly evolving sites of the GBP1 PBM are sufficient to abolish or restore bacterial detection abilities, illustrating a role for epistasis in the evolution of pathogen recognition. We further demonstrate that the squirrel monkey GBP2 C-terminal domain recently gained the ability to target *S. flexneri* through a stepwise process of convergent evolution. These findings reveal a mechanism by which accelerated evolution of a PBM shifts GBP target specificity and aid in resolving the molecular basis of GBP function in cell-autonomous immune defense.

## Introduction

Diverse metazoan cell types possess the innate ability to resist infection by pathogens, a feature termed cell-autonomous immunity. Detection of intracellular bacteria, viruses, or eukaryotic parasites by host factors engenders cell-autonomous defense programs operating to contain or eliminate invasive pathogens from an infected cell (1). These defense programs can be activated in response to proinflammatory interferons produced by professional immune or neighboring infected cells. Interferon signaling prompts the expression of interferon-stimulated genes (ISGs) which encode a wide range of antimicrobial proteins (2). Among the most highly-upregulated ISGs are a class of dynamin-related cytoplasmic GTPases called guanylate binding proteins, or GBPs. Vertebrate GBPs contribute to defense against diverse pathogens, and GBP function has also been implicated in the regulation of inflammation (3–5).

GBPs consist of an N-terminal catalytic GTPase domain followed by an elongated helical domain which mediates interactions with target proteins or membranes. GTP binding and hydrolysis promotes the dimerization, oligomerization, and polymerization of GBPs as well as recruitment of additional GBP family members (6). Oligomerization of GBPs on pathogen-containing membrane-bound compartments prompts an array of antimicrobial activities including the production of radical oxygen species by co-recruited oxidases, the fusion of these compartments with degradative lysosomes, their encapsulation within autophagosome-like structures, or the lytic disintegration of microbe-containing compartments (7). Some GBPs also possess the ability to target microbes that reside inside the host cell cytosol. Cytosolic bacteria enclosed by GBPs undergo lytic destruction in mouse macrophages (8–10) or are blocked from engaging the host actin polymerization machinery in human epithelial cells, thereby losing the ability to disseminate (11, 12). The importance of GBPs as potent immune effectors is further illustrated by the recent discovery that the enteric bacterial pathogen *Shigella flexneri* injects host cells with the virulence factor IpaH9.8 which specifically disrupts GBP function (11–13). IpaH9.8 is an E3 ubiquitin ligase that directly binds to several GBP family members, targeting them for destruction by the proteasome (12–14). While IpaH9.8 is so far the only reported microbial GBP antagonist, it is likely that additional pathogen-encoded GBP countermeasures remain to be discovered.

Despite a wealth of evidence supporting the role of GBPs in cell-autonomous host defense, the molecular mechanisms underlying GBP function and target specificity remain enigmatic. One informative observation is that mammalian GBPs target cytosolic microbes as well as microbe-associated membranous structures in a hierarchical manner, with individual GBPs functioning as ‘pioneers’ that recruit other family members through heterotypic interactions (15, 16). In particular, GBP1, GBP2, and GBP5 in humans are predicted to directly associate with target membranes due to the presence of a C-terminal CaaX box leading to post-translational prenylation, which acts as a hydrophobic lipid anchor (6). In support of this model, it was shown that recombinant human GBP1 (hGBP1) binds directly to lipid bilayers *in vitro* in a GTP- and prenylation-dependent manner (17). However, prenylation alone is unlikely to provide targeting specificity and other protein motifs are expected to enable prenylated hGBPs to discriminate between ‘self’ and ‘non-self’ membranes inside infected cells (18). Consistent with this hypothesis, we previously demonstrated that hGBP1 is unique amongst all human GBPs in its ability to target cytosolic *S. flexneri* due to the presence of a polybasic motif (PBM) positioned immediately adjacent to its C-terminal CaaX box (11). We further provided genetic evidence for a possible functional interaction between the PBM of hGBP1 and the O-antigen moiety of lipopolysaccharide (LPS), the major component of the Gram-negative bacterial envelope, implying the so far untested model that the PBM of hGBP1 physically engages LPS O-antigen in order to bind to the surface of *S. flexneri* and likely other Gram-negative bacteria (11).

The unique ability of hGBP1 among all human GBPs to target cytosolic Gram-negative bacteria, the expansion of the *GBP* gene family in humans and other species as well as the diversity of targets recognized by distinct GBP isoforms suggests a model in which individual GBPs have evolved unique characteristics to recognize and respond to pathogens spanning the entire tree of life. It is also notable that mouse Gbp2, the closest murine homolog of hGBP1, lacks a clearly defined C-terminal PBM and yet is capable of recognizing and eliminating cytosolic *S. flexneri (13)*, indicating some variability in the molecular interactions that promote bacterial detection by GBPs. Collectively, these findings suggest that the divergence of GBPs within and between host genomes has drastically shifted bacterial recognition function, potentially in response to antagonistic coevolution with pathogens. In the current study we set out to address this question focusing on a subset of primate GBPs which possess the ability to specifically recognize and bind cytosolic bacteria. While genetic variation in GBPs is likely to alter recognition of various microbes, we focused our investigation on *S. flexneri* as a model cytosolic Gram-negative bacterium whose virulence is strongly diminished by GBP recruitment. Moreover, we anticipate that variation in cytoplasmic bacterial surfaces could have provided a potent selective force for GBP adaptation across vertebrates. Through a combination of phylogenetic and experimental approaches, we find that accelerated evolution of membrane-targeting motifs in GBP1 and GBP2 have led to repeated gain, loss, and enhancement of bacterial detection abilities in primates.

## Results

### Divergence and evolution of prenylated GBPs in simian primates

We chose to focus our initial investigation on the prenylated GBPs (GBP1, GBP2, and GBP5, Fig. 1A) which are predicted to directly interact with intracellular microbes or microbe-derived membranous structures such as bacterial outer membrane vesicles (19, 20). While human GBP1, GBP2, and GBP5 all possess the CaaX motif required for post-translational prenylation, GBP1 alone possesses a PBM which contributes to cytoplasmic bacterial recognition (Fig. 1B). We first noted a large-scale genomic deletion encompassing the *GBP5* locus in several Old World monkeys, suggesting that GBP5 is absent in this family (Fig. 1C). Alignment of GBP1 and GBP2 amino acid sequences from simian primates revealed another surprising observation. While the C-terminal CaaX box is highly conserved among GBP1 and GBP2 orthologs, the amino acid sequence immediately adjacent exhibits an extreme degree of amino acid divergence (Fig. S1). Notably, this region encompasses the C-terminal PBM of hGBP1, a protein motif essential for the hGBP1-mediated recognition of cytosolic *S. flexneri* in human epithelial cells (11). We considered why a domain that is required for pathogen recognition might be subject to such extreme sequence variation, despite strict conservation of the CaaX box. One possible explanation for this divergence is that prenylation of the GBP1 and GBP2 CaaX box confers general membrane-anchoring properties, while the adjacent C-terminal amino acid sequences allow these GBPs to discriminate between microbial ‘non-self’ and ‘self’ membrane surfaces. Rapid diversification of GBP1 and GBP2 orthologs in this case suggests the existence of repeated evolutionary conflicts between cytoplasmic pathogens and GBPs, in which pathogen alterations to membrane surface molecules mediates evasion of GBP targeting. Our initial observations revealing elevated genetic diversity in the C-terminal regions of primate GBP1 and GBP2 thus mandated further evolutionary and experimental investigation.

**Figure 1.**
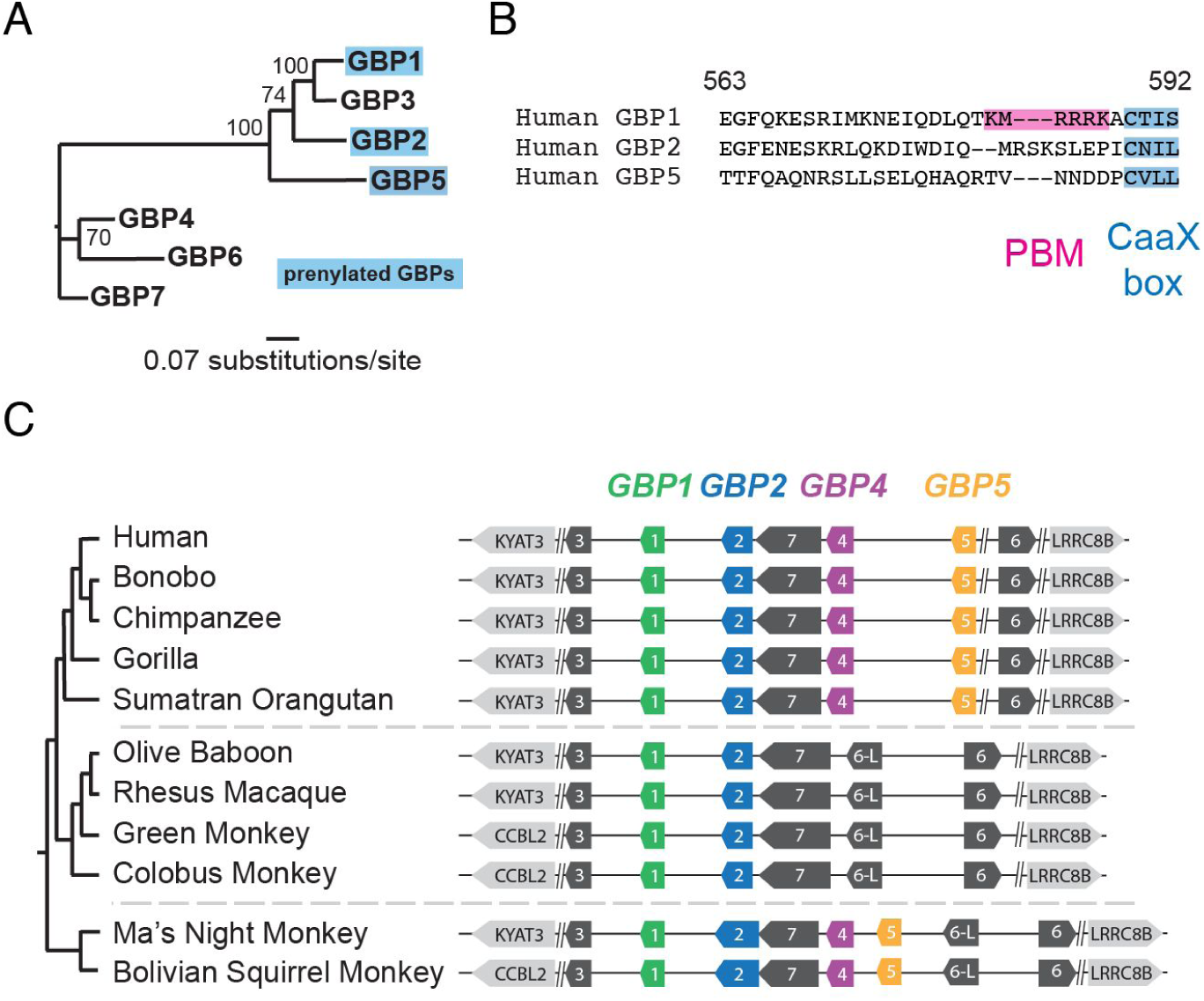
Diversity and evolution of primate guanylate binding proteins. **A.** Maximum-likelihood phylogeny of the seven human GBP family members. GBPs which undergo post-translational prenylation are highlighted. Bootstrap values at tree nodes are based on 1000 replicates. **B**. Amino acid alignment of the C-terminal region of human GBP1, GBP2, and GBP5. The location of the CaaX motif, which undergoes prenylation, as well as the polybasic motif (PBM) of GBP1 are highlighted. Amino acid numbering is relative to human GBP1. **C**. Diagram of the GBP gene cluster from representative primate genomes, illustrating the apparent loss of GBP4 and GBP5 in Old World monkeys.

### Rapid diversification of primate GBP1 and GBP2 C-terminal domains

If GBP C-terminal genetic variants provide a fitness advantage to the host in the face of pathogen antagonism, evolutionary theory predicts such variants could rapidly and repeatedly spread through host populations due to the forces of positive selection (also termed diversifying selection). One method to infer instances of repeated positive selection in protein coding genes is through calculation of the ratio of nonsynonymous substitutions per nonsynonymous site relative to synonymous substitutions per synonymous site, referred to as dN/dS or ω. An elevated dN/dS ratio greater than 1 indicates that amino acid substitutions have fixed in populations more rapidly than expected by chance, consistent with positive selection acting preferentially on beneficial mutations (21). To detect potential signatures of positive selection in GBP1 and GBP2, we compiled gene orthologs from simian primates by direct Sanger sequencing of cDNA from primate cell lines as well as from the Genbank database (Fig. 2A). We then subjected GBP1 and GBP2 datasets to phylogenetic tests estimating dN/dS at individual sites, implemented through the PAML and HyPhy software packages (22, 23) (see Materials and Methods). All tests identified statistically significant support for positive selection acting on both GBP1 (Fig. 2B, S1, Tables S1-S4) and GBP2 (Fig. 2C, S2, Tables S4-S8). Notably, multiple positions in the C-terminal regions of both GBP1 and GBP2 exhibit signatures of positive selection, whereas the adjacent CaaX box is highly conserved. We observed that the highest degree of variation in these sites appears to be found in New World primates, which diverged from the common ancestor of humans roughly 40 million years ago. These findings suggest that both GBP1 and GBP2 have been subject to repeated positive selection in the primate lineage, including at sites in the PBM which promote intracellular pathogen recognition.

**Figure 2.**
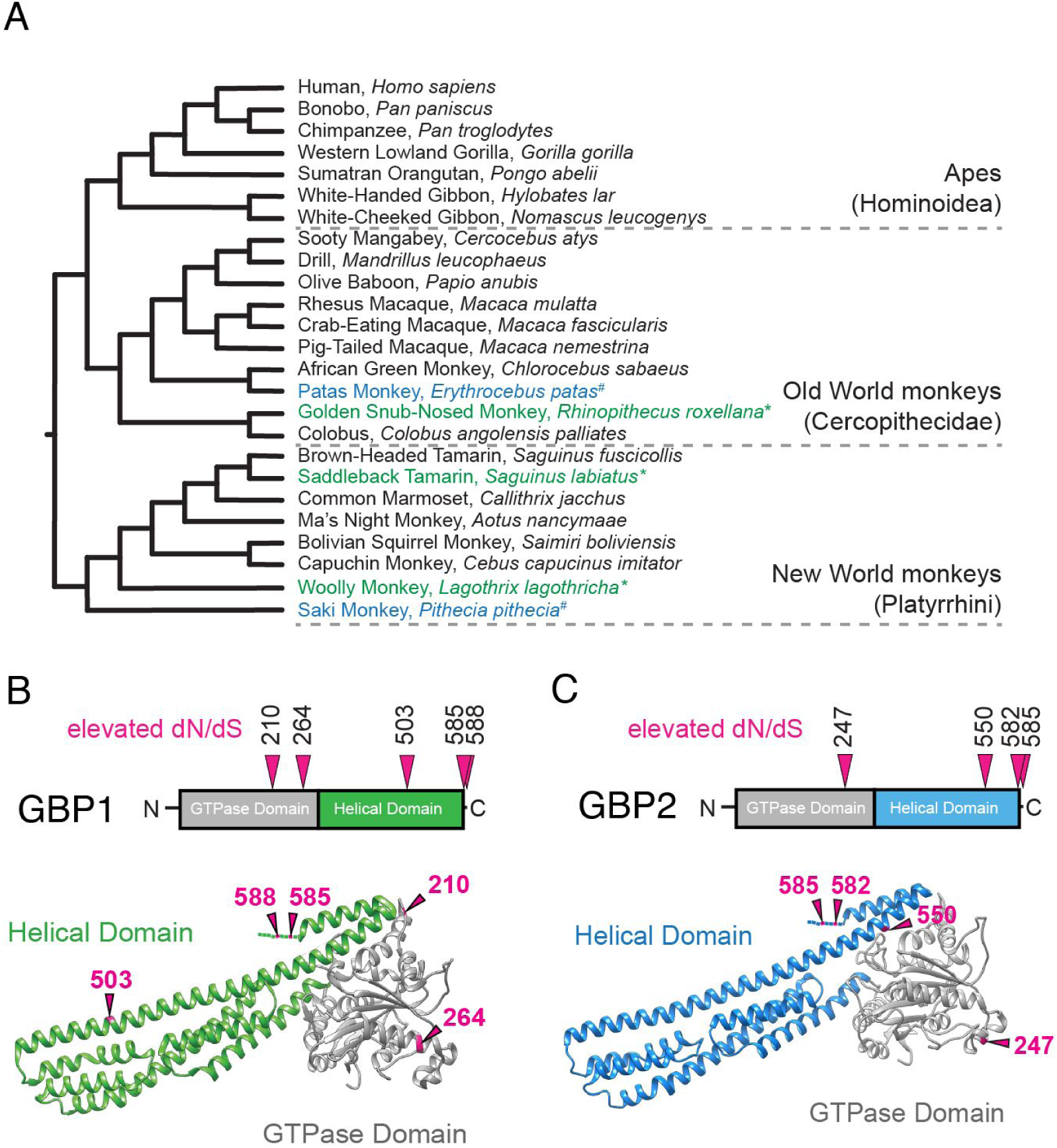
Accelerated evolution of the C-terminal polybasic motif of primate GBP1 and GBP2. **A**. Species tree of simian primates used for phylogenetic analyses. Species in black were included in both the GBP1 and GBP2 datasets. * Indicates species that were included in GBP1 analysis alone. # Indicates species that were included in GBP2 analysis alone. **B**. Sites in GBP1 that display a statistically significant elevation in dN/dS ratio, indicative of repeated positive selection. Site numbers correspond to human GBP1 and were identified using at least four separate inference methods in the PAML and HyPhy software packages. **C**. Sites in GBP2 that display a statistically significant elevation in dN/dS ratio, indicative of repeated positive selection, Site numbers correspond to human GBP2 and were detected using at least four separate inference methods as in B.

### The variable polybasic motif of primate GBP1 modulates targeting of pathogenic *Shigella flexneri*

We next sought to determine how rapid divergence in the PBM (Fig. 3A) impacts pathogen recognition function of GBP1. We generated a series of protein chimeras in which the C-terminal PBM of hGBP1 (576 QDLQTKMRRRKACTIS 592) was replaced with the orthologous sequence from other primate GBP1 alleles (Fig. 3B). Human GBP1 as well as the chimeric constructs were expressed using an anhydrotetracycline (aTc)-inducible system in CRISPR-engineered hGBP1-deficient HeLa cells (*GBP1*^KO^) to ensure that any targeting activity observed was due to variation in exogenously-expressed GBPs. To assess the consequences of GBP1 function, we chose *S. flexneri* as a model pathogen given it is targeted specifically by hGBP1 and its dissemination within the host is highly sensitive to GBP recruitment (11–13). Cells were infected with GFP-expressing wildtype *S. flexneri* strain 2457T or the coisogenic *ΔipaH9.8* mutant. Consistent with previous results (11–13), we found that hGBP1 targeting to bacteria was dependent on the triple arginine stretch of its PBM and blocked by the *S. flexneri* hGBP1 antagonist IpaH9.8 (Fig. S3). To avoid confounding results related to IpaH9.8 antagonism of GBPs, we conducted all subsequent experiments comparing the targeting efficiencies of GBP variants using the *ΔipaH9.8* mutant. For our initial studies we generated chimeras using C-terminal domains from a single representative hominoid (white-cheeked gibbon, *Nomascus leucogenys*), Old World monkey (rhesus macaque, *Macaca mulatta*), and New World monkey (Ma’s night monkey, *Aotus nancymaae*) as well as the triple-arginine PBM mutant hGBP1^R584-586A^ as a negative control. Performing these experiments with chimeric proteins allowed us to control for interspecific sequence differences outside the C-terminal region of GBP1. These experiments revealed that despite significant sequence divergence, the C-terminal domains of gibbon, rhesus macaque, and night monkey were all capable of targeting cytosolic *S. flexneri* (Fig. 3C). In fact, we observed that the night monkey GBP1 C-terminal amino acid stretch possesses significantly enhanced targeting ability relative to hGBP1 (Fig. 3D). aTc-induced protein expression levels were comparable across all GBP1 chimeras and mutants in the absence or presence of *S. flexneri* infections (Fig. 3E, S4), suggesting that the enhanced targeting of the night monkey C-terminus was a result of specific amino acid substitutions. To further explore the consequences of GBP1 diversity in other New World primates, we generated additional chimeras using squirrel monkey (*Saimiri boliviensis*), capuchin (*Cebus capucinus imitator*), and marmoset (*Callithrix jacchus*) GBP1. Both squirrel monkey and capuchin GBP1 C-terminal motifs also displayed improved GBP1 targeting relative to human (Fig. 3F). In contrast, the marmoset GBP1 chimeric protein poorly associated with cytosolic *S. flexneri* (Figs. 3F-G). This reduced function could be the result of an insertion of a stretch of five neutral, mostly hydrophobic amino acids (NVFFP) into the PBM of marmoset GBP1, which is not present in other primates (Fig. 3A). Collectively, these results demonstrate that the ability to target intracytosolic *S. flexneri* has been enhanced and lost in distinct lineages of New World primates.

**Figure 3.**
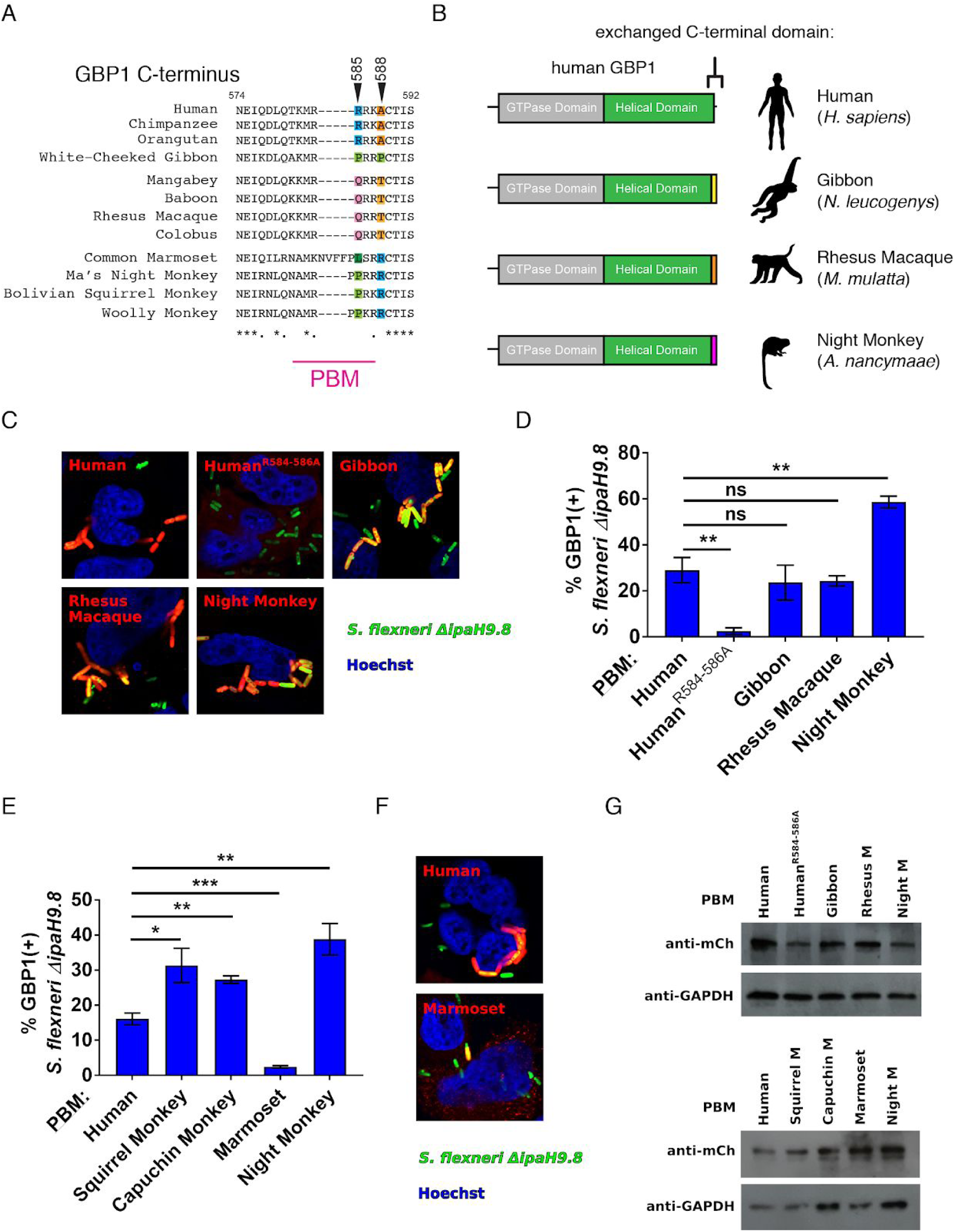
Diversification of the GBP1 polybasic motif in primates enhances recognition of *Shigella flexneri*. **A**. Alignment of the C-terminal region of GBP1 across several simian primates. Positions that were identified with significantly elevated dN/dS are indicated with arrows. The area corresponding to the PBM in human GBP1 is marked below. **B**. Diagram illustrating gene chimeras in which the human GBP1 C-terminus was replaced with corresponding sequences from related primate species. These mCherry-tagged constructs were expressed in *GBP1*^*KO*^ Hela cells for subsequent experiments. **C**. Representative immunofluorescence images of *GBP*1^*KO*^ cells expressing GBP1 chimeras (red) infected with GFP-expressing *S. flexneri ΔipaH9.8* strain (green). Hoechst stain of DNA is shown in blue. D. Quantification of intracellular *S. flexneri* co-localizing with mCherry GBP1 as in C. Bar graphs show means ±SEM from three independent experiments. **E**. Quantification of GBP1-*S. flexneri* co-localization using New World monkey GBP1 chimera constructs. Bar graphs show means ±SEM from four independent experiments. **F**. Representative immunofluorescence images comparing hGBPl and hGBP1-marmoset chimeric constructs, as in F. Significance was determined by unpaired two-tailed t-tests relative to results for human GBP1. * p ≦ 0.05; **, p ≦ 0.01; ***, p ≦ 0.001; ns, nonsignificant. **G**. Western blot from GBP1 chimera-expressing cells. mCherry antibody was used to detect individual GBP1 protein expression. GAPDH expression was included as a loading control.

### Genetic interactions constrain evolution of the GBP1 polybasic motif

To gain a more detailed understanding of how natural selection has shaped the evolution of GBP1 function, we focused on two positions corresponding to R585 and A588 in hGBP1 that exhibit signatures of repeated positive selection across primates (Fig. 4A). We initially hypothesized that mutating each position in human GBP1 to the corresponding amino acid in night monkey GBP1 may be sufficient to confer improved bacterial recognition activity. We introduced single amino acid substitutions at both sites in hGBP1 to amino acids found in night monkey GBP1, as well as generating a double mutant protein. In contrast to our early expectations, substitution of arginine at position 585 to proline significantly abrogated human GBP1 binding to *S. flexneri*. Substituting alanine 588 to arginine did not enhance targeting, but introducing the R585P mutation into this background preserved targeting to *S. flexneri* (Fig. 4B-C). These results illustrate that, despite their rapid divergence, intramolecular epistasis between sites in the PBM constrains available evolutionary trajectories that maintain antibacterial function. Our findings also indicate that there are additional sequence features in the New World monkey PBM beyond positions 585 and 588 that must contribute to its enhanced bacterial targeting relative to hGBP1.

**Figure 4.**
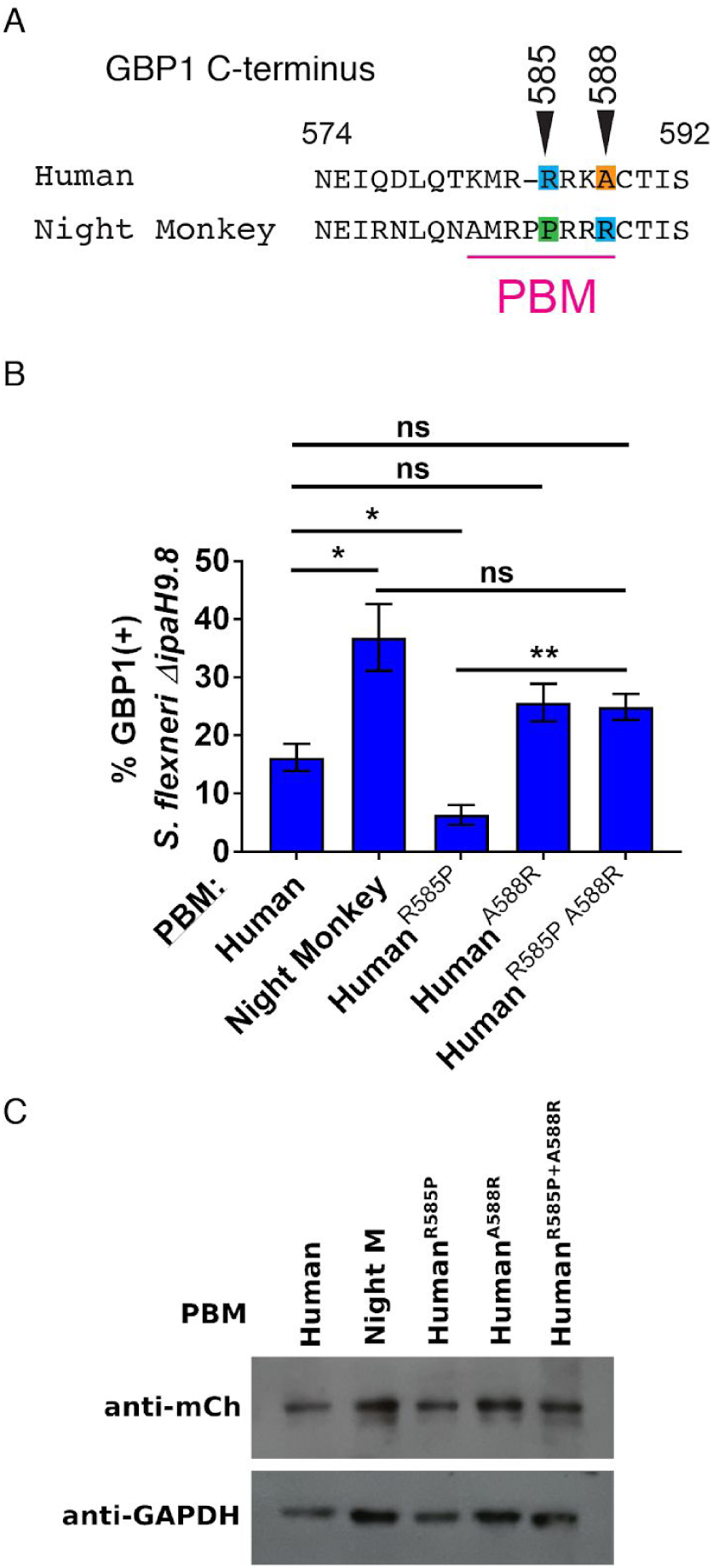
Epistasis among rapidly-diverging sites constrains bacterial recognition by GBP1. **A.** Amino acid alignment of the human and night monkey GBP1 C-terminal regions. Sites 585 and 588 which exhibit signatures of positive selection across primates are indicated. **B**. Quantification of GBP1-S. *flexneri* co-localization using hGBP1, the hGBP1-night monkey chimera, as well as single and double point mutations at sites 585 and 588 in human GBP1. Bar graphs show means ±SEM from three independent experiments. **C**. Western blot from GBP1-expressing cells. mCherry antibody was used to detect individual GBP1 protein expression. GAPDH expression was included as a loading control. Significance was determined by unpaired two-tailed t-tests relative to results for human GBP1 or as indicated. *, p ≦ 0.05; **, p ≦ 0.01; ns, nonsignificant.

### Convergent evolution of bacterial recognition by squirrel monkey GBP2

Similar to hGBP1, human GBP2 (hGBP2) undergoes prenylation via its conserved C-terminal CaaX box. hGBP2 can co-localize with cytosolic *S. flexneri* through heterotypic interactions with hGBP1, but fails to target *S. flexneri* in *GBP1*^KO^ cells due to the lack of an appropriate C-terminal targeting motif (11–13). Our earlier phylogenetic analyses revealed that prenylated GBP paralogs are highly divergent at the unstructured C-terminal region immediately preceding the CaaX box, suggesting that unique C-terminal sequences direct individual prenylated hGBP isoforms towards distinct microbial targets. According to this model, we expect that the C-terminal residues of prenylated GBPs could be subject to conflict with intracellular pathogens evolving to evade recognition. This is consistent with the high degree of divergence amongst the C-termini of GBP2 in primates, suggesting that its putative antimicrobial targeting specificity may have undergone shifts during recent primate evolution. In particular, we observed that the C-terminus of GBP2 in squirrel monkeys contains a series of substitutions as well as a small deletion which produce in a sequence that closely resembles the GBP1 PBM (Fig. 5A). This sequence was observed both in the publically available Bolivian squirrel monkey (*Saimiri boliviensis*) genome as well as confirmed by direct Sanger sequencing of *GBP2* from the related common squirrel monkey (*Saimiri scruitius*). The corresponding sequence of GBP2 from capuchin monkeys, a close relative of squirrel monkeys, was highly divergent suggesting most of these alterations arose recently in the *Saimiri* lineage (Fig. 5A). To determine if the squirrel monkey GBP2 PBM is sufficient to promote targeting of intracellular bacteria, we replaced the C-terminal region of human GBP1 with that of squirrel monkey GBP2. We observed that the resulting GBP1-GBP2 chimeric protein was able to localize to intracellular *S. flexneri*, albeit at lower frequencies than hGBP1. This finding indicates that squirrel monkey GBP2 gained the ability to target *S. flexneri* and potentially other intracellular Gram-negative bacteria through an example of recent convergent evolution.

**Figure 5.**
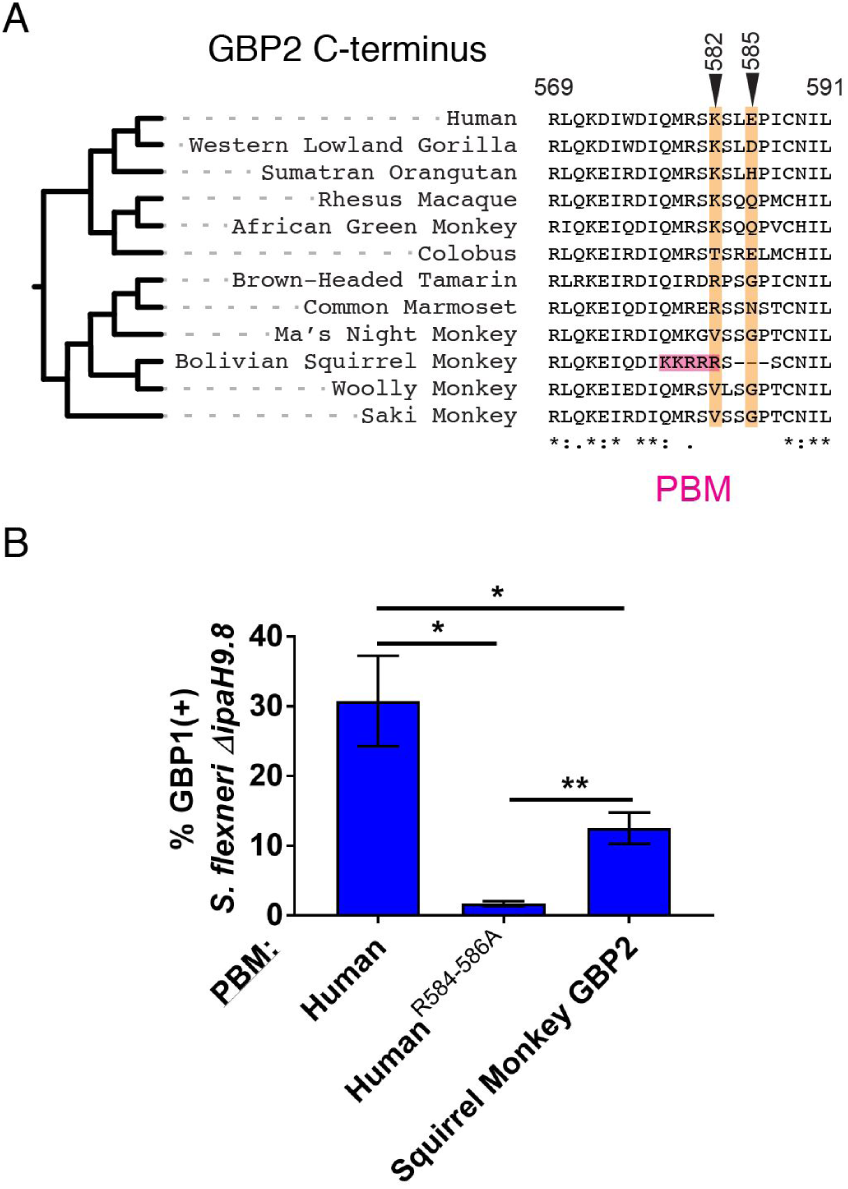
Convergent evolution of bacterial recognition by a squirrel monkey GBP2 polybasic motif. **A.** Amino acid alignment of the C-terminal regions of primate GBP2 orthologs. Sites exhibiting signatures of positive selection across species are denoted in orange. Pink region highlights the emergence of a polybasic motif in squirrel monkey GBP2. **B**. Quantification of GBP-S. *flexneri* co-localization using hGBP1, the hGBP1-PBM mutant, as well as a chimeric hGBP1 fused to the C-terminal region of squirrel monkey GBP2. Bar graphs show means±SEM from four independent experiments. Significance was determined by unpaired two-tailed t-tests relative to results for human GBP1 or as indicated. *,p ≦ 0.05; **, p ≦ 0.01.

## Discussion

GBPs continue to emerge as critical mediators of vertebrate cell-autonomous immunity, contributing to resistance against diverse pathogens as well as susceptibility to inflammatory disease. Although GBPs exhibit variation in both gene copy number and amino acid identity within and between species, the consequences of such genetic variation for GBP function have remained unclear. The present study illustrates how rapid evolution of the C-terminal PBM in GBP1 and GBP2 controls detection of a host cytosol-invading bacterial pathogen in humans and related simian primates. These findings are consistent with a model in which beneficial PBM mutations that enhance pathogen recognition have rapidly spread through host populations by the process of positive selection. The observed patterns of repeated selection in GBP1 and GBP2 could reflect two scenarios, the first of which being a classic ‘arms race’ in which GBPs and specific bacterial surface molecules (such as LPS O-antigens or porin proteins) antagonistically co-evolve to improve or impair recognition of bacterial surfaces, respectively. These patterns could also arise from fluctuations in the types of pathogens that have impose selection on host populations over time, with PBM mutations mediating alterations in the spectrum of targets recognized by a single GBP. This second case is analogous to selection in a fluctuating environment, rather than strict co-evolution. It is likely that both scenarios have influenced the GBP family during vertebrate evolution. While our studies have focused on host-*S. flexneri* interactions as a model system, we expect that PBM variation has impacted GBP activity towards a range of cytosolic bacterial pathogens during the course of vertebrate evolution. The diversification of GBP pathogen targeting capabilities is also highly relevant to animal models of infection as the mouse ortholog of hGBP1, mouse GBP2, detects cytosolic *S. flexneri* through a poorly-defined process (13) independent of a bona-fide PBM (24). Given the dynamic changes observed in a subset of simian primates, it is likely that GBPs from other non-model vertebrates could harbor as yet undiscovered pathogen targeting capabilities.

The evolution-guided experimental approach applied here provides new details regarding the molecular basis of GBP target recognition. The high degree of conservation in GBP1 and GBP2 CaaX boxes suggests that post-translational prenylation and subsequent membrane association has been critical for the function of both proteins. These findings agree with numerous studies illustrating how these GBPs are able to associate with diverse intracellular membranes that include pathogen-containing vacuoles, such as those occupied by the bacterium *Chlamydia* or the protist *Toxoplasma*, viral replication complexes and bacterial cell envelopes (5, 25, 26). By first associating with these target membranes, GBP1 and GBP2 are able to recruit other GBP family members as well as additional immune effectors such as the immunity related GTPases (27). In contrast to CaaX box conservation, dynamic evolution of the adjacent PBM is indicative of selective pressures to target rapidly diversifying pathogen targets. Our results suggest that the PBM could function as an intracellular ‘zip code,’ allowing GBPs to distinguish between foreign or ‘non-self’ and ‘self’ membrane surfaces (18). In this respect it is also of note that the CaaX box and PBM are present in a wide range of GTPases that do not perform primary roles in cell-autonomous immunity, including Rab and Rho GTPases (28). It is thus tempting to speculate that ancestral interferon-stimulated GTPase function may have evolved from a more promiscuous prenylated GTPase capable of executing intracellular housekeeping functions. This model is consistent with our recent finding that hGBP1 as well as mouse GBPs are able to detect vacuolar membrane damage and to intersect with the galectin protein family involved in the removal of damaged organelles (29). Future evolution-guided molecular studies could aid in understanding the types of PBM-substrate interactions that underlie the diversity of cellular functions that depend on CaaX-proximal PBMs. In this regard, the evolution of the GBP PBM resembles a model of evolutionary “tinkering” proposed by François Jacob (30) and reported in other cases of protein diversification (31). Among closely related primates we observe instances of enhanced targeting ability in New World monkeys, but also cases where new mutations have attenuated GBP function as with marmoset GBP1. Mutation of individual sites in the PBM of GBP1 further demonstrates that epistasis could strongly constrain evolutionary paths to improved or novel functions. The fact that vertebrate genomes often encode several GBP family members with cooperative and overlapping targeting abilities may relax selective constraint on single GBP genes to allow for this exploration of broader sequence space. GBPs therefore provide an attractive and tractable model to investigate fundamental questions on evolutionary novelty and innovation.

Much work delineating the molecular mechanisms of host-microbe genetic conflict has focused on interactions with viruses (32–38), although emerging studies from ourselves and others highlight the potential for pathogenic bacteria to promote similar evolutionary dynamics (39–41). Given the ability of GBPs to target a diverse array of pathogens and pathogen-containing compartments, future studies aimed at understanding potential tradeoffs in target specificity during GBP evolution would greatly improve our understanding of their functions in cell-autonomous immunity. The ability of GBPs to also cooperate and form heteromeric complexes likely further serves to enhance their breadth in pathogen recognition.

In addition to investigations of GBP family evolution reported here, previous work has established that members of the myxovirus resistance (Mx) protein family of interferon-stimulated GTPases has also been subject to repeated positive selection in primates (42, 43) as well as counter-adaptation by viral pathogens (44). While GBP and Mx protein family GTPases differ in their molecular targets and the specific mechanism by which pathogen recognition occurs, they may share fundamental principles underlying their immune surveillance functions. Mammalian Mx protein diversity, particularly within the L4 loop of the alpha helical stalk region, controls the breadth and specificity of viral proteins recognized by this restriction factor (42, 45, 46). The combination of Mx protein oligomerization and L4 loop flexibility could provide a broad target interface to mediate the interaction of these antiviral GTPase with diverse viral protein substrates (47). Similar to the L4 loop of Mx proteins, we propose that the unstructured C-terminal regions preceding the CaaX boxes of GBP1, GBP2, and GBP5 confer target specificities and equip these prenylated proteins with the ability to associate with pathogen membranes or pathogen-containing membrane-bound compartments. Parallels between the evolution of the PBM of GBPs and the L4 loop of MxA are indicative of diverse intracellular pathogens exerting selective pressure on both protein families in different host species. Recent studies of MxA diversity further highlight the potential for tradeoffs between breadth and specificity of antiviral activity during the evolution of the L4 loop (48). Such observations are consistent with both dynamic changes in copy number and sequence variation of interferon-stimulated GTPases in order to target diverse pathogens.

Recent studies have indicated that *S. flexneri* encodes a secreted effector protein, IpaH9.8, which targets GBPs for degradation by the proteasome (12, 13). Although our preliminary studies did not reveal any significant differences in the ability of IpaH9.8 to antagonize GBP variants, it is entirely possible that other microbial GBP inhibitors have also imposed selective pressure on this gene family during animal evolution. In support of this hypothesis, our phylogenetic analyses identified signatures of positive selection acting on sites beyond the PBM, namely in the GTPase and alpha-helical domains of GBP1 and GBP2 (Fig. 2). Future studies may resolve if and how GBP evolution impacts resistance to other as yet to be discovered pathogen-encoded inhibitors. Together this study establishes functional links between GBP evolution and the molecular basis of intracellular bacterial pathogen recognition.

## Materials & Methods

### Primate GBP Genetic Sources

Primate *GBP1* and *GBP2* sequences were retrieved from NCBI GenBank entries for primates with sequenced genomes. For other primates, sequences were obtained by Sanger sequencing of PCR amplicons using cDNA isolated from cell lines obtained from Coriell Cell Repositories (Camden, NJ). Briefly, RNA was harvested using the ZR-*Duet*™ DNA/RNA MiniPrep Plus kit (Zymo Research). Isolated RNA (50ug) from cell lines was used as a template for RT-PCR (SuperScript III; Invitrogen). Sequences of interest were PCR amplified from cDNA using Phusion High-Fidelity mastermix (Thermo) according to the manufacturer’s protocol and were cloned in to pCR2.1 (Invitrogen). Sanger sequencing was performed from at least three individual clones. *GBP1* and *GBP2* gene sequences obtained from the NCBI database included human (*Homo sapiens*), chimpanzee (*Pan troglodytes*), pygmy chimpanzee (*Pan paniscus*), Western lowland gorilla (*Gorilla gorilla*), Sumatran orangutan (*Pongo abelii*), sooty mangaby (*Cercocebus atys*), drill (*Mandrillus leucophaeus*), olive baboon (*Papio anubis*), Rhesus macaque (*Macaca mulatta*), crab-eating macaque (*Macaca fascicularis*), pit-tailed macaque (*Macaca nemestrina*), green monkey (*Chlorocebus sabaeus*), colobus (*Colobus angolensis palliates*), common marmoset (*Callithrix jacchus*), Ma’s night monkey (*Aotus nancymaae*), capuchin monkey (*Cebus capucinus imitator*), and Bolivian squirrel monkey (*Saimiri boliviensis*). The *GBP1* orthologs cloned from cDNA (with Coriell Identifier [ID] numbers) are as follows: white-handed gibbon (PR01121), white-cheeked gibbon (PR00712), red-chested mustached tamarin (AG05308), saddleback tamarin (AG05313), common woolly monkey (AG05356). The *GBP2* orthologs cloned from cDNA (with Coriell Identifier [ID] numbers) are as follows: patas monkey (AG06116), red-chested mustached tamarin (AG05308), common squirrel monkey (AG05311), common woolly monkey (AG05356), white-faced saki (PR00239). GBP gene sequence data from this project has been deposited in GenBank.

### GBP Phylogenetic and Protein Structure Analysis

DNA multiple sequence alignments were performed using MUSCLE and indels were manually edited based on amino-acid comparisons. Phylogenetic trees for each sequence set were derived from generally accepted primate relationships (49). Maximum-likelihood analysis of the *GBP1* and *GBP2* data sets were performed with codeml of the PAML software package (22). Positive selection was assessed by fitting the multiple alignment to either F3X4 or F61 codon frequency models. Likelihood ratio tests (LRTs) were performed by comparing the following site-specific models (NS sites): M1 (neutral) with M2 (selection), and M7 (neutral, beta distribution of dN/dS<1) with M8 (selection, beta distribution, dN/dS>1 allowed). PAML analysis identified sets of amino acids with high posterior probabilities (more than 0.95) for positive selection by a Bayesian approach. Additional LRTs from the HyPhy software package which account for synonymous rate variation and recombination (FEL, SLAC, MEME) were performed using the Datamonkey server (23). Sites under positive selection for GBP1 were mapped onto three-dimensional molecular structures available from the Protein Data Bank (PDB ID 1DG3) using Chimera (50) (http://www.cglu.ucsf.edu/chimera/). GBP2 sites of positive selection were mapped onto a three-dimensional molecular structure generated using the I-Tasser modeling program provided by the University of Michigan (https://zhanglab.ccmb.med.umich.edu/I-TASSER).

### Design of GBP expression constructs

Plasmids encoding mCherry-tagged hGBP1 and a triple arginine mutation in the hGBP1 PBM were previously reported (11). The mCherry-tagged hGBP1 plasmids was used as a template to generate hGBP1-primate GBP chimeras. First, a BglII restriction site within the linker-sequence separating the N-terminal mCherry-tag from hGBP1 was eliminated in pmCherry-hGBP1 by Quickchange Site Directed Mutagenesis (Agilent) using the oligomer pair pmCherry-hGBP1DBglII-F and -R (Table S9). Next, 5’-Kozak-mCherry-hGBP1 from the resulting vector were amplified with oligomers that simultaneously added 5’-attB1 and 3’-attB2 sites, introduced a BglII site spanning hGBP1 codons Q577-L579 via a synonymous mutation in codon Q577 (CA<u>G</u> to CA<u>A</u>), and truncated GBP1 beyond codon L579 (attB1-mCherry-F and attB2-hGBP1DC_BglII-R, Table S9). This PCR product was inserted into pDONR221 (Invitrogen) via Gateway BP recombination (Invitrogen). Sequences encoding primate PBMs were added to the resulting pDONR221-mCherry-human GBP1DC_BglII vector following BglII digestion using Ligation-Independent cloning (In-Fusion, Clontech) with annealed oligomers that also restored human GBP1 Q580-L581 and the human GBP1 CaaX box (Table S9). Finally, resulting chimeras were inserted into the lentiviral Tetracycline-inducible vector pInducer20 (51) by Gateway LR recombination (Invitrogen).

5’-Kozak-mCherry-hGBP1 was cloned into pDONR221 by Gateway BP recombination following amplification with primers attB1-mCherry-F and attB2-hGBP1-R, followed by insertion into pInducer20 by Gateway LR recombination. Mutant R585P and A588R alleles were constructed from pDONR221-mCherry-human GBP1 by Quickchange Site Directed Mutagenesis using oligomer pairs hGBP1_R585P-F and -R, hGBP1_A588R-F and -R, and hGBP1_R585P_A588R-F and -R (Table S9). Resulting mutant constructs were inserted into pInducer20 via Gateway LR recombination.

To construct the chimera in which the C-terminal portion of hGBP1 is replaced with that of Bolivian Squirrel Monkey GBP2, a derivative of pmCherry-human GBP1 was used in which a synonymous mutation was made within the flexible region separating alpha helices 11 and 12 to introduce a BclI restriction enzyme site. This plasmid was propagated in *dam/dcm*^-^*E. coli* (New England Biolabs), and a synthetic “gBlock” encoding Bolivian Squirrel Monkey GBP2 residues 476-588 was inserted via BclI restriction digest/ligation.

### Cell lines, cell culture, and ectopic gene expression

hGBP1-deficient HeLa cells (*GBP1*^KO^) were described previously (11). Unless noted otherwise, *GBP1*^KO^ cells were stably transduced with an (aTc)-inducible gene expression systems to drive the expression of hGBP1 as well as its mutant and chimera variants. For transient transfection experiments cells were transfected with indicated expression constructs using Lipofectamin LTX (Thermofisher Scientific). Cells were cultivated in Dulbecco’s Modified Eagle Medium (Gibco, Thermo Fisher Scientific) supplemented with 10% Fetal Bovine Serum (Corning), 1% non-essential amino acids (Sigma), and 55 µM β-mercaptoethanol (Gibco, Thermo Fisher Scientific) at 37 °C and 5% CO_2_.

### Bacterial strains and infections

*GBP1*^*KO*^ Hela cells were cultured on glass coverslides and infected with GFP-expressing *S. flexneri* strain 2547T or the coisogenic D*ipaH9.8* GFP^+^ mutant strain at an MOI of 50, essentially as described (11). Briefly, tryptic soy broth (TSB) supplemented with 50 µg/ml carbenicillin was inoculated with a single Congo red-positive colony and grown overnight at 37°C with shaking. Stationary overnight cultures were diluted 1:30 in 5 ml of fresh TSB and incubated for 1 h to 1.5 h at 37°C with shaking until an OD_600_ of 0.4-0.6 was reached. Bacteria were diluted in prewarmed cell culture medium and spun onto host cells for 10 min at 700 × *g*. Infected cells were incubated for 30 min at 37°C and 5% CO_2_ and subsequently washed twice with Hanks balanced salt solution (HBSS), followed by addition of cell culture medium containing 25 mg/ml gentamicin. Cells were incubated for an additional 2.5 h at 37°C and 5% and then fixed in 4% paraformaldehyde for 15 min at room temperature and mounted onto glass slides for fluorescence microscopy. Fixed cells were imaged using an Axio Observer.Z1 microscope (Zeiss) and image analysis to quantify co-localizaiton of mCherry fusion proteins with bacteria was performed as described previously (11).

## Acknowledgements

We are grateful to members of the Barber and Coers laboratories for critical reading of the manuscript. This work was supported by National Institutes of Health grants R00GM115822 (to M.F.B.), AI103197, and AI139425 (to J.C.). M.F.B. holds a Faculty Scholar Award from the Donald E. and Delia B. Baxter Foundation. J.C. holds an Investigator in the Pathogenesis of Infectious Disease Award from the Burroughs Wellcome Fund.

