## Supplemental Figures & Tables for "A rapidly evolving polybasic motif modulates bacterial detection by guanylate binding proteins"

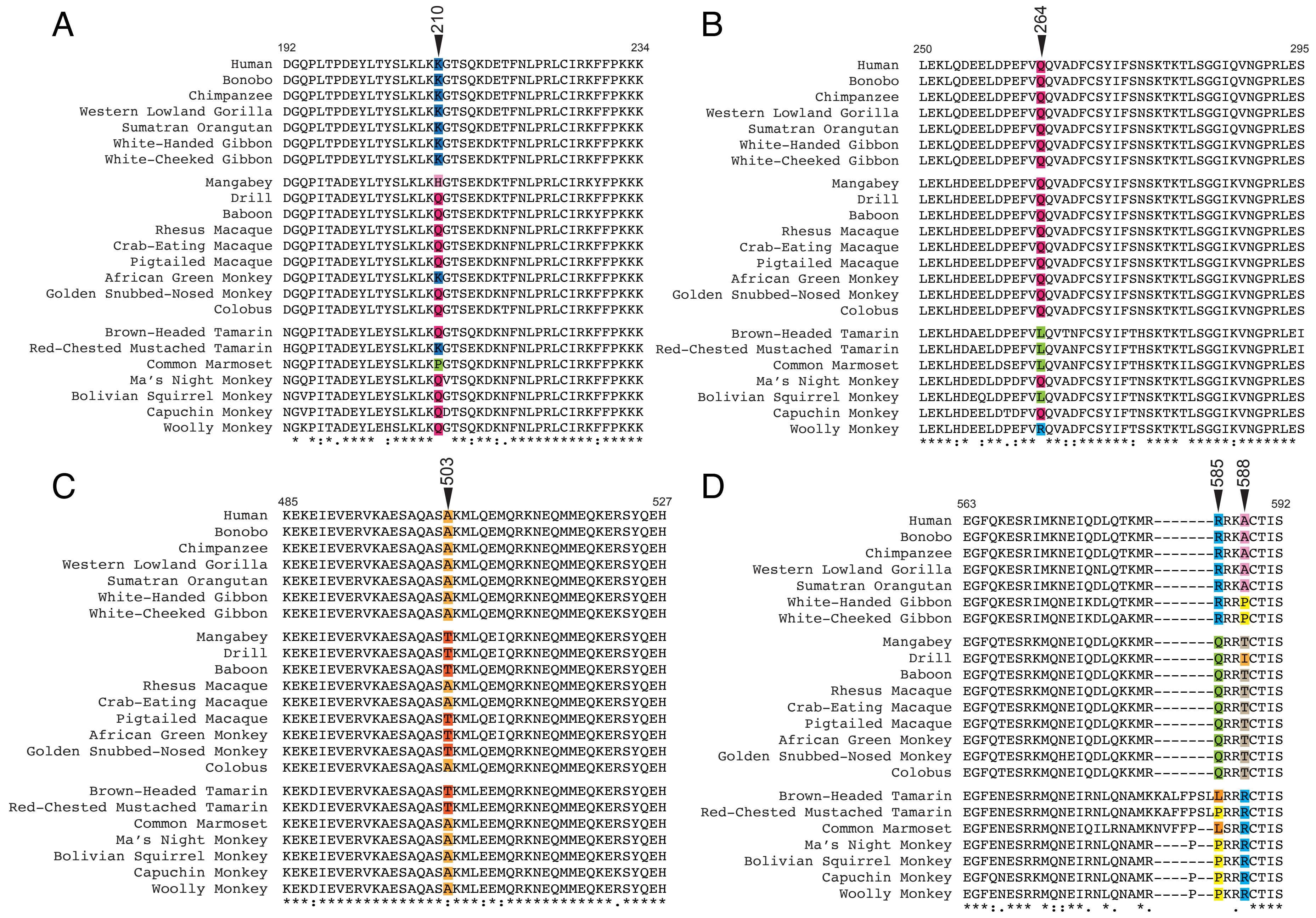

**Figure S1.** Amino acid alignments of GBP1 orthologs surrounding sites of elevated dN/dS as identified by PAML and HyPhy.

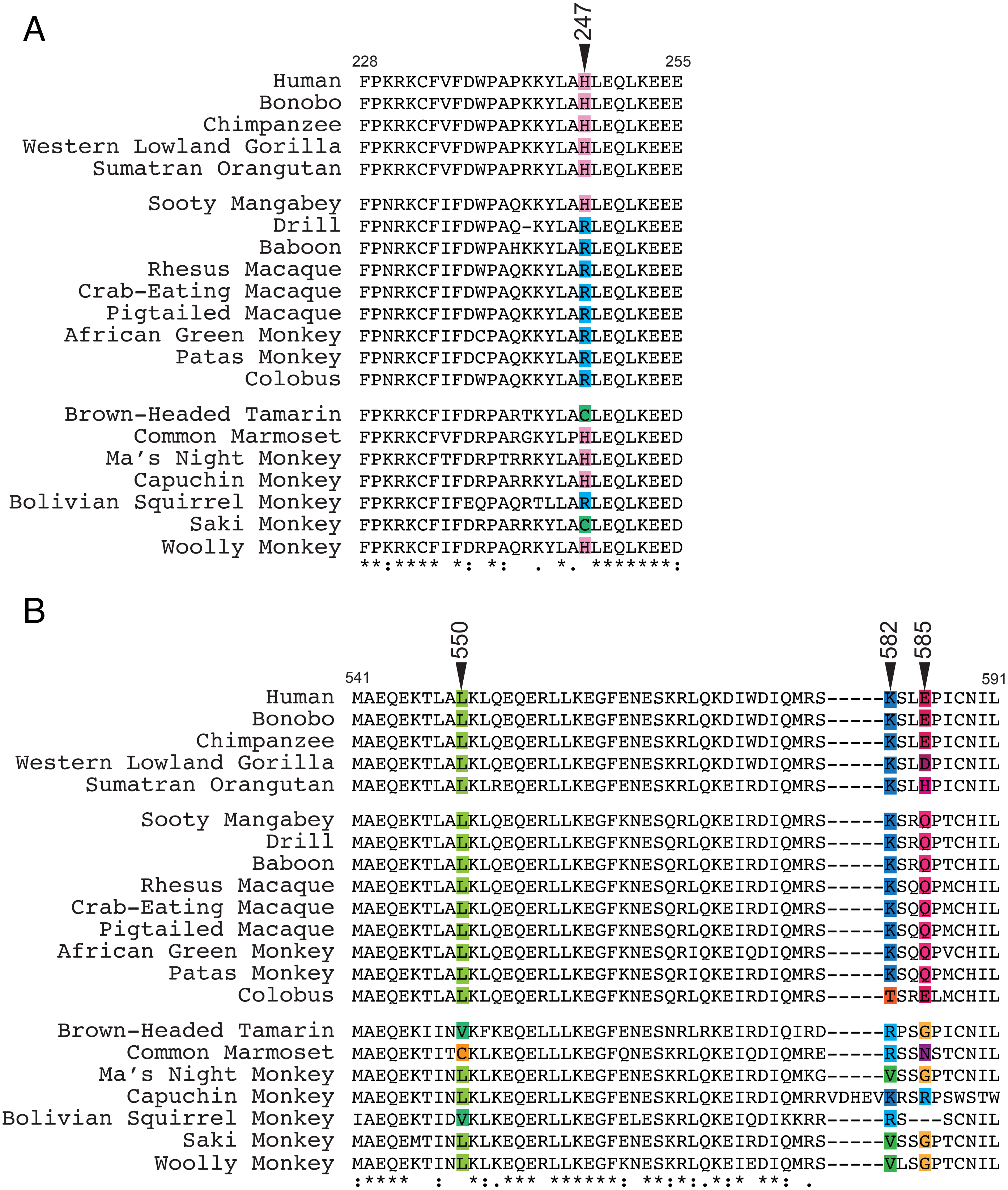


**Figure S2.** Amino acid alignments of GBP2 orthologs surrounding sites of elevated dN/dS as identified by PAML and HyPhy.

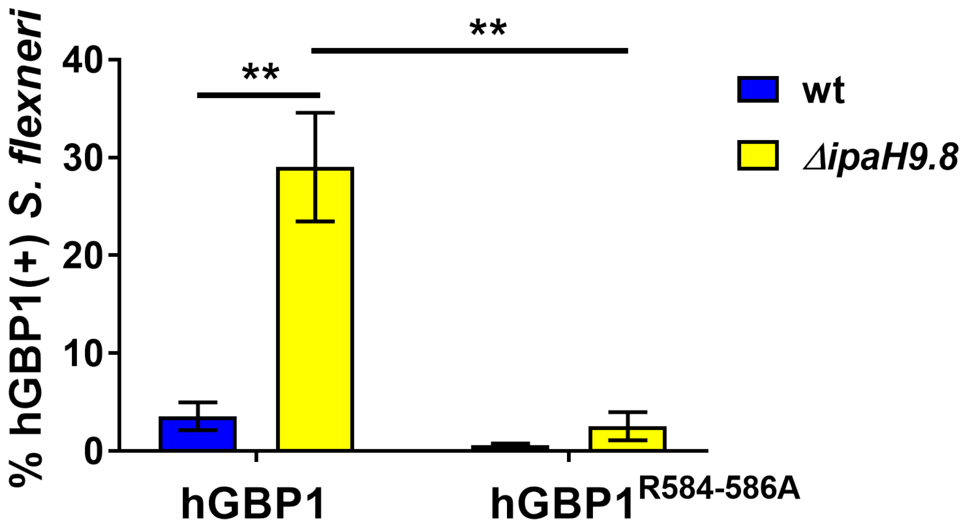


**Figure S3.** Quantification of intracellular *S. flexneri* co-localizing with mCherry GBP1. Experiments were performed using wildtype *S. flexneri* or a mutant strain containing a deletion of the IpaH9.8 effector which targets GBP1 for degradation. Bar graphs show means ±SEM from three independent experiments.

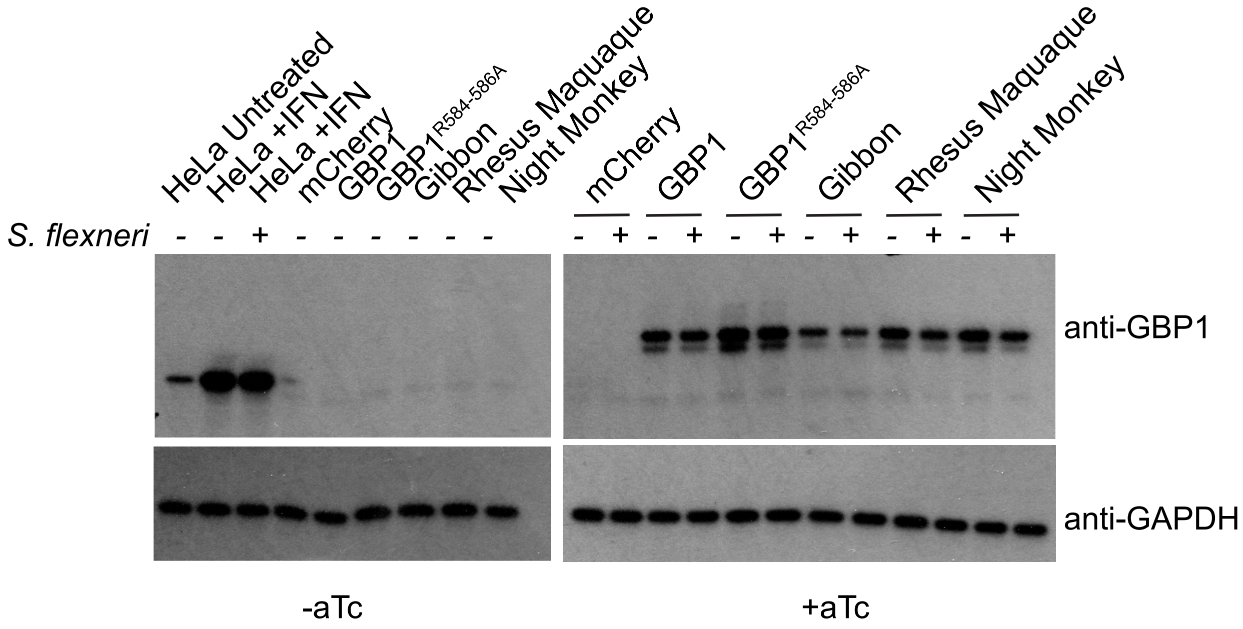


**Figure S4.** HeLa or GBP1-KO HeLa cells complemented with pInducer-mCherry-GBP1 constructs were stimulated overnight with either 200 U/ml IFNgamma (HeLa +IFNg controls) or 0.5 ug/ml aTc. The following day, cells were left uninfected or infected with WT *S. flexneri* at an MOI of 50 for 3 h. Western blots detecting GBP1 or GAPDH loading control are indicated.

**Table S1.** GBP1 whole gene log likelihood scores and parameter estimates for four models of variable $\omega$ among sites assuming the f3X4 model of codon frequencies (PAML).

| Site Model | Parameter Estimates | Sites* with $\omega$**>1 | lnL |
|---|---|---|---|
| **M1: Neutral** | ($\omega_0$=0) $f_0$=0.624<br>($\omega_1$=1) $f_1$=0.376<br>branch $\omega$ (mean)=0.376 | Not allowed | -4986.95 |
| **M2: Selection** | ($\omega_0$=0) $f_0$=0.614<br>($\omega_1$=1) $f_1$=0.374<br>**($\omega_2$=5.70) $f_2$=0.012**<br>branch $\omega$ (mean)=0.440 | 448Y 0.955 | -4981.31 |
| **M7: $\beta$** | $p$=0.00500<br>$q$=0.00750<br>branch $\omega$ (mean)=0.400 | Not allowed | -4987.21 |
| **M8: $\beta$ and $\omega$** | $p$=0.01644 $q$= 0.02948<br>$f_0$=0.987<br>**$\omega_1$=5.41 ( $f_1$=0.013)**<br>branch $\omega$ (mean)=0.432 | **210 K** 0.964<br>424 A 0.953<br>448 Y 0.986<br>**585 R** 0.976 | -4981.27 |

*posterior probabilities >0.95 by Bayes Empirical Bayes (BEB) analysis
**$\omega$=dN/dS
***Amino acid positions shown are for human GBP1.

**Table S2.** Summary of positive selection in primate GBP1 (MEME, FEL, SLAC).

| Model | Sites with evidence of positive selection (p-value)* |
|---|---|
| MEME | 194 Q 0.013<br>203 T 0.059<br>**210 K 0.028**<br>249 Q 0.038<br>**264 Q 0.064**<br>**503 A 0.071**<br>562 K 0.067<br>582 K 0.055<br>**585R 0.035**<br>586R 0.034<br>**588A 0.088** |
| FEL | **210 K 0.022**<br>**264 Q 0.045**<br>**503 A 0.051**<br>**585 R 0.025**<br>**588 A 0.066** |
| SLAC | No positively selected sites identified |

*Amino acid positions shown are for human GBP1.

**Table S3.** Summary of positive selection in primate GBP1 using REL algorithm.

| Sites with evidence of diversifying selection* | Posterior probability β>α |
|---|---|
| Y143 | 0.990 |
| H150 | 0.991 |
| E218 | 0.990 |
| D192 | 0.990 |
| Q194 | 0.979 |
| T203 | 0.973 |
| **K210** | **0.998** |
| G211 | 0.992 |
| Q214 | 0.977 |
| T218 | 0.997 |
| F229 | 0.988 |
| A248 | 0.975 |
| Q249 | 0.993 |
| E257 | 0.991 |
| P260 | 0.974 |
| E261 | 0.993 |
| **Q264** | **0.999** |
| I332 | 0.993 |
| T349 | 0.993 |
| D359 | 0.989 |
| E363 | 0.992 |
| E389 | 0.994 |
| D405 | 0.991 |
| A409 | 0.993 |
| V413 | 0.991 |
| A424 | 0.965 |
| Y447 | 0.990 |
| I455 | 0.980 |
| T463 | 0.985 |
| E484 | 0.994 |
| **A503** | **0.999** |
| M509 | 0.965 |
| R522 | 0.991 |
| N537 | 0.997 |
| V540 | 0.973 |
| Q559 | 0.996 |
| Q566 | 0.994 |
| K567 | 0.991 |
| I571 | 0.997 |
| Q577 | 0.998 |
| L578 | 0.995 |
| K582 | 0.994 |
| R584 | 0.973 |
| **R585** | **0.995** |
| R586 | 0.992 |
| K587 | 0.991 |
| **A588** | **0.999** |

*Amino acid positions shown are for human GBP1.

**Table S4.** Summary of positive selection in primate GBP1 using FUBAR algorithm.

| Sites with evidence of diversifying selection* | Posterior probability $\beta>\alpha$ |
|---|---|
| **K210** | **0.987** |
| **Q264** | **0.921** |
| **A503** | **9.932** |
| **R585** | **0.981** |
| **A588** | **0.929** |

*Amino acid positions shown are for human GBP1.

**Table S5.** GBP2 whole gene log likelihood scores and parameter estimates for four models of variable $\omega$ among sites assuming the f3X4 model of codon frequencies (PAML).

| Site Model | Parameter Estimates | Sites* with $\omega$**>1 | lnL |
|---|---|---|---|
| **M1: Neutral** | ($\omega_0$=0) $f_0$=0.735<br>($\omega_1$=1) $f_1$=0.265<br>branch $\omega$ (mean)=0.293 | Not allowed | -5061.10 |
| **M2: Selection** | ($\omega_0$=0) $f_0$=0.732<br>($\omega_1$=1) $f_1$=0.248<br>**($\omega_2$=6.6) $f_2$=0.021**<br>branch $\omega$ (mean)=0.421 | **247 H 0.993**<br>575 W 0.976<br>**582 K 0.999**<br>**585 E 1.000**<br>587 I 0.975 | -5037.85 |
| **M7: $\beta$** | $p$=0.01204 $q$=0.02377<br>branch $\omega$ (mean)=0.314 | Not allowed | -5061.77 |
| **M8: $\beta$ and $\omega$** | $p$=0.07563 $q$=0.18279<br>$f_0$=0.97860<br>**$\omega_1$=6.5 ($f_1$=0.0214)**<br>branch $\omega$ (mean)=0.426 | 241 P 0.969<br>**247 H 0.997**<br>575 W 0.991<br>**582 K 0.999**<br>583 S 0.979<br>584 L 0.978<br>**585 E 1.000**<br>587 I 0.993 | -5038.30 |

*posterior probabilities >0.95 by Bayes Empirical Bayes (BEB) analysis
**$\omega$=dN/dS
***Amino acid positions shown are for human GBP2.

**Table S6.** Summary of positive selection in primate GBP2 (MEME, FEL, SLAC).

| **Model** | **Sites with evidence of positive selection (p-value)*** |
|---|---|
| MEME | 208 K (0.04)<br>209 G (0.04)<br>**247 H (0.03)**<br>296 L (0.01)<br>513 E (0.01)<br>565 N (0.02)<br>**582 K (0.01)**<br>**585 E (0.04)**<br>586 P (0.01) |
| FEL | **247 H (0.050)**<br>**582 K (0.033)**<br>**585 E (0.033)** |
| SLAC | No positively selected sites identified |

*Amino acid positions shown are for human GBP2.

**Table S7.** Summary of positive selection in primate GBP2 using REL algorithm.

| Sites with evidence of diversifying selection* | Posterior probability $\beta > \alpha$ |
|---|---|
| H247 | 0.98 |
| K582 | 0.98 |
| E585 | 0.99 |

*Amino acid positions shown are for human GBP2.

**Table S8.** Summary of positive selection in primate GBP2 using FUBAR algorithm.

| **Sites with evidence of diversifying selection*** | **Posterior probability $\beta>\alpha$** |
|---|---|
| M11 | 0.930 |
| N161 | 0.925 |
| E188 | 0.944 |
| F234 | 0.926 |
| A241 | 0.940 |
| **H247** | **0.984** |
| E327 | 0.939 |
| E334 | 0.924 |
| G338 | 0.999 |
| A549 | 0.909 |
| **L550** | **0.936** |
| E564 | 0.905 |
| **K582** | **0.989** |
| **E585** | **0.992** |

*Amino acid positions shown are for human GBP2.

**Table S9.** Oligonucleotides used in this study.

| Oligo Name | Sequence |
|---|---|
| pmCherry-hGBP1DBglII-F | 5'-GCTGTACAAGTCCGGACTCAGGTCCATGGCATCAGAGATC-3' |
| pmCherry-hGBP1DBglII-R | 5'-GATCTCTGATGCCATGGACCTGAGTCCGGACTTGTACAGC-3' |
| attB1-mCherry-F | 5'-GGGGACAAGTTTGTACAAAAAAGCAGGCTGCCACCATGGTGAGCAAGGGCGAGG-3' |
| attB2-hGBP1DC_BglII-R | 5'-GGGGACCACTTTGTACAAGAAAGCTGGGTCTTAGAGATCTTGTATCTCATTTTTCATT ATTCTGCTTTC-3' |
| Gibbon_PBM-F | 5'-ATGAGATACAAGATCTCCAGGCGAAAATGCGTCCGCGCCGTCCGTGTACCATAAGCTAAGATCTCTAAGACCCA-3' |
| Gibbon_PBM-R | 5'-TGGGTCTTAGAGATCTTAGCTTATGGTACACGGACGGCGCGGACGCATTTTCGCCTGGAGATCTTGTATCTCAT-3' |
| Rhesus Macaque_PBM-F | 5'-ATGAGATACAAGATCTCCAGAAGAAAATGCGTCAGCGCCGTACCTGTACCATAAGCTAAGATCTCTAAGACCCA-3' |
| Rhesus Macaque_PBM-R | 5'-TGGGTCTTAGAGATCTTAGCTTATGGTACAGGTACGGCGCTGACGCATTTTCTTCTGGAGATCTTGTATCTCAT-3' |
| Night Monkey_PBM-F | 5'-ATGAGATACAAGATCTCCAGAACGCGATGCGTCCGCCGCGTCGCCGTTGTACCATAAGCTAAGATCTCTAAGACCCA-3' |
| Night Monkey_PBM-R | 5'-TGGGTCTTAGAGATCTTAGCTTATGGTACAACGGCGACGCGGCGGACGCATCGCGTTCTGGAGATCTTGTATCTCAT-3' |
| Capuchin_PBM-F | 5'-ATGAGATACAAGATCTCCAGAACGCCATGAAGCCCCCTCGGCGCAGATGTACCATAAGCTAAGATCTCTAAGACCCA-3' |
| Capuchin_PBM-R | 5'-TGGGTCTTAGAGATCTTAGCTTATGGTACATCTGCGCCGAGGGGGCTTCATGGCGTTCTGGAGATCTTGTATCTCAT-3' |
| Squirrel Monkey_PBM-F | 5'-ATGAGATACAAGATCTCCAGAACGCCATGAGACCCCGCAAGAGATGTACCATAAGCTAAGATCTCTAAGACCCA-3' |
| Squirrel Monkey_PBM-R | 5'-TGGGTCTTAGAGATCTTAGCTTATGGTACATCTCTTGCGGGGTCTCATGGCGTTCTGGAGATCTTGTATCTCAT-3' |
| Marmoset_PBM-F | 5'-ATGAGATACAAGATCTCCAGAACGCCATGAAGAACGTGTTCTTCCCCCTGTCCCGGCGCTGTACCATAAGCTAAGATCTCTAAGACCCA-3' |
| Marmoset_PBM-R | 5'-TGGGTCTTAGAGATCTTAGCTTATGGTACAGCGCCGGGACAGGGGGAAGAACACGTTCTTCATGGCGTTCTGGAGATCTTGTATCTCAT-3' |
| attB2-hGBP1-R | 5'-GGGGACCACTTTGTACAAGAAAGCTGGGTCTTAGCTTATGGTACATGCCTTTCGTC-3' |
| hGBP1_R585P-F | 5'-CAGACGAAAATGAGACCACGAAAGGCATGTACCATAAGC-3' |
| hGBP1_R585P-R | 5'-GCTTATGGTACATGCCTTTCGTGGTCTCATTTTCGTCTG-3' |
| hGBP1_A588R-F | 5'-CAGACGAAAATGAGACGACGAAAGCGATGTACCATAAGC-3 |
| hGBP1_A588R-R | 5'-GCTTATGGTACATCGCTTTCGTCGTCTCATTTTCGTCTG-3' |
| hGBP1_R585P_A588R-F | 5'-CTCCAGACGAAAATGAGACCACGAAAGCGATGTACCATAAGC-3' |
| hGBP1_R585P_A588R-R | 5'-GCTTATGGTACATCGCTTTCGTGGTCTCATTTTCGTCTGGAG-3' |
| Squirrel Monkey GBP2 C-Terminus gBlock | 5'-CTTCTACAAACTGATCAGTCACTCTCAGAAAAGGAAAAAGCGCTTGAAGTGGAACGTGTAAAGGCTGAATCTGCCGAAGCTGCAAAGAAAATGTTGGAGGAAATACAAAAGAAGAACCAGCAGATGATGGAACAGAAAGAGAAGAGTTATCAGGAACATGTGAAACAATTGACTGAGAAGATGGAGAGTGATAGGGCCCAATTAATAGCGGAGCAAGAGAAGACCATCGATGTTAAACTTAAGGAACAGGAACGCCTTCTCAAAGAGGGATTCGAGCTTGAGAGCAAGAGACTTCAAAAAGAGATACAAGATATCAAGAAAAGACGCAGATCATCATGTAACATACTCTAATGATCATAATCAGCCA-3' |
